# Virtual World Coupling with Photosynthesis Evaluation for Synthetic Data Production

**DOI:** 10.1101/2025.02.06.633870

**Authors:** Dirk N. Baker, Mona Giraud, Jens Henrik Göbbert, Hanno Scharr, Morris Riedel, Ebba Þóra Hvannberg, Andrea Schnepf

## Abstract

In this work, we couple the functional-structural plant model CPlantBox to the Unreal Engine by exploiting the implemented raytracing pipeline to evaluate light influx on the plant surface. There are many approaches for photosynthesis computation and light evaluation, though they typically are limited by versatility, compute speed, or operate on much coarser resolutions. This work specifically addresses the concern that data generation pipelines tend to be unresponsive and do not include model-based knowledge as part of the generation pipeline. Using established photosynthesis solvers, we model the interaction between the Unreal Engine and the FSPM to measure physical properties in the virtual world. This is successful if we are able to reproduce experimental results using an in silico model. As part of the pipeline, we generate a surface geometry and utilize material shaders that are designed to establish a baseline surface model for light interception and transmission, based on simple parameter sets that can be calibrated. Using a Selhausen field experiment as baseline, we reproduce the photosynthesis effectiveness of the plants in the 2016 winter wheat experiments. Our pipeline is deeply intertwined with data generation and has been proven to perform well at scale. In this work, we build on our previous work by showcasing both a simulation study of a light evaluation as well as quantifying how well our system performs on high-performance computing systems.

## 1 Introduction

Simulating the real world is a pathway to deeper understanding of its mechanisms. This is especially true in plant science, where many individually distinct processes jointly form emergent behavior of a plant (Thornley 2011; Walia et al. 2024). Here, studying the influences of the environment, or of plant properties, becomes crucial to fully explore crop behavior as agriculture becomes more sustainable and resilient. Having access to a model of the real world, particularly in cases for which reference experimental data is available but sparse, can assist with filling gaps in the data. A use-case for this is to map the experimental results onto synthetic data for deep learning (DL) models. Particularly in plant sciences, where image analysis is a critical tool and the primary bottleneck (Minervini et al. 2015; Taiz 2013; Tsaftaris et al. 2016), using model-based data sources can uncover hidden mechanisms. A faithful version of an experiment would include a calibrated structure of a plant, producing both images and dense training labels. The potential to uncover systematic image features that plants with a certain trait exhibit, is the first and most important step towards training models that detect such traits remotely. Embedding models into the data generation pipeline is especially impactful as a common critique of data production pipelines is their unresponsiveness and one-directional character, e.g. as reported by Li et al. (2024). In this work, we showcase a method to embed and communicate with the virtual world in a way that does not only produce more useful data labels for a concrete problem such as photosynthesis, but also explore the expressiveness of our virtual world model to accommodate these models. Functional-structural plant models (FSPMs), such as CPlantBox (Giraud et al. 2023) seldomly represent only one functional process and are instead a coupling of different models (Sievänen et al. 2014). Plant structural definitions can inform boundaries of plant processes, such as leaf surface, or root-shoot-interface, where e.g. flux parameters can be defined that are communicated between the organs. A key feature, however, is the fact that the plant structure is incorporated explicitly rather than using heuristics to signify plant behavior. Though heuristics can be established using plant simulations, particularly for upscaling as investigated by Vanderborght et al. (2024), plant simulations are typically performed to gain insights into plant mechanisms.

Photosynthetic simulations with stomatal regulation generally solve a fixed-point iteration (Von Caemerer 2013) and the reactions involved in the uptake of co_2_ are solved on the basis of a steady reaction flow limited by availability of nutrients (Giraud et al. 2023) or stomatal metabolism (Yin et al. 2004). The CPlantBox module for photosynthesis, developed by Giraud et al. (2023), combines different models (Farquhar et al. 1980; Leuning 1995; Tuzet et al. 2003) into a whole-plant framework that couples the water and carbon dynamics from root to leaf. Although the model can use specific input data for each leaf section, Giraud et al. (2023) only used mean atmospheric data as basic heuristical approach. Models that calculate the actual light influx in a plant involve radiative transfer modeling (RTM), which describes the light propagation and eventual impact onto the surface of the plant. RTM has standardized concepts (John V. Martonchik and Strahler 2000; Nicodemus et al. 1977) and is implemented in a variety of models, particularly DART (J. P. Gastellu-Etchegorry and Gascon 2004). For the retrieval of functional properties from remote sensing data, empirical models such as LESS developed by Qi et al. (2019) and extended e.g. by Zhao et al. (2024), have been successfully implemented. Another example of a high-quality model for radiative transfer available as open-source is prospect, developed by Féret and Boissieu (2024). Worth noting is that similar concepts and models have been implemented when a rendering technique called *physics-based rendering* was established, and the similarity between the two fields regarding this specific use-case is of particular note, as described by Salesin et al. (2024).

In experiments to parameterize FSPMs, particularly the measurement of the structural portion can yield much information on the performance of the crop. Diversity of structure, e.g. in root systems, has been identified to be one of the primary indicators of adaptation of a crop to its environment by Yu et al. (2024). As such, many data generation pipelines make use of a plant’s structure to accommodate variability (Bailey 2019; Baker et al. 2023; Ward et al. 2018). The key factor for the embedding of an FSPM for data production is a reasonable reproduction of the target space, which is RGB images for many applications. In our pipeline, we are using the Unreal Engine (UE) to compute the virtual world. UE is a multimedia 3D graphics engine, which has seen use across industries, including its origin in gaming, simulation science (Agarwal et al. 2023), plant science (Li et al. 2024), and virtual reality (Krüger et al. 2024). Our own recent work brought forth the Synavis framework, a library designed to dynamically setup, steer, and measure within the UE virtual world (Baker et al. 2023, 2024).

An embedding recently implemented in the HELIOS pipeline by Lei et al. (2024) shows the potential for pipelines to accommodate functional information. We believe that the inclusion of an FSPM that can simulate functions in the whole plant (roots and shoot) and at a sub-organ level provides a valuable extension to this by allowing the models to communicate between domains (soil - plant - atmosphere) and scales (plant section - field). Thus, in this work, we extend the Synavis framework to work together with the photosynthesis module of CPlantBox, which until recently employed a uniform light influx heuristic. Our implementation of this pipeline uses the physics-based rendering employed in UE to simulate light exposure to achieve an accurate light exposure computation for CPlantBox, which we validate by modeling a digital twin of an experiment at the Field Minirhizotron Facility in Selhausen. The data-based pipeline that is enabled in Synavis is extended by using a model-view which takes physical properties to allow radiative transfer modelling and a data view that is able to map functional photosynthesis data onto a synthetic data pipeline, which we showcase by the example of light exposure efficiency. In our replication, we make use of experimental data provided by the TERENO platform (Bogena 2016; Reichenau et al. 2020) to re-create a field experiment while also retaining a high level of structural and functional information through the use of a full-scale plant model. Lastly, our pipeline is HPC compatible, with a focus on ensuring that parallelization is feasible, and we include full scaling experiments with this manuscript.

## 2 Methods

We will be highlighting selected aspects of the FSPM along with the geometric embedding into the virtual scene as part of our solution to compute light influx. Note that for certain parts, this paper includes expressions and terms that are otherwise written as render code, and for further resources, we need to refer to the open-source implementation of UE^1^.

### 2.1 Virtual World Generation in Synavis

We used the Synavis framework to create a virtual world in UE. An in-depth description of the Synavis framework can be found in our recent paper (Baker et al. 2023). Synavis is a framework for the coupling of simulations with virtual environments, allowing the sampling and measuring of values through the framework for the purposes of training neural networks. In this work, we extended this functionality by allowing the FSPM simulation to access data from the virtual environment directly, allowing the superimposition of plant function onto the virtual environment and the subsequent measurement physical properties in the environment. Within UE, we used the base rendering framework as well as the entity-component system implemented in UE. We developed a framework to measure light intensity, a pipeline to embed model data into UE to generate synthetic data based on FSPM simulation results, as well as a pipeline to setup and manage distributed virtual scene handling through Synavis.

Our setup has three distinct aspects: the CPlantBox simulation, the UE model scene, and the resulting visualization. These aspects of the setup can be separated and communicated asynchronously, but in our setups we typically assume that each application is running concurrently (Baker et al. 2024). The field setup configures and runs the CPlantBox simulations. The visualizer carries vertex-triangle information that can be used to map functional properties. Through Synavis, plant geometries are inserted into the virtual scene once every time step. A texture containing surface properties is being generated per organ because of how the texture wraps around the plant, see Fig. 1 and Sec.2.2. Light influx measurement is done after thsse radiation calibration, submitted as point measurement command to the `ALightMeter` class in UE, which carries out the measurement. Necessary parameters are meter resolution, measurement delay to accommodate scene update and measurement duration to accommodate path tracing. Path tracing is a UE term that refers to the physics-based raytracing of image pixels without heuristic. UE does implement a heuristic for raytracing, called Lumen, which uses a reduced-detail scene to complete the light information and might not correctly reflect the physical properties of the surfaces, as reported by Agarwal et al. (2023) in a remote sensing context. The measurement is passed back through Synavis as influx per segment, which is passed to the Photosynthesis module. The parametrization is entirely on the Python side, which retains all functional parameters and values. Mapping of the photosynthesis output is done exactly like the mapping of functional properties.

**Fig. 1:**
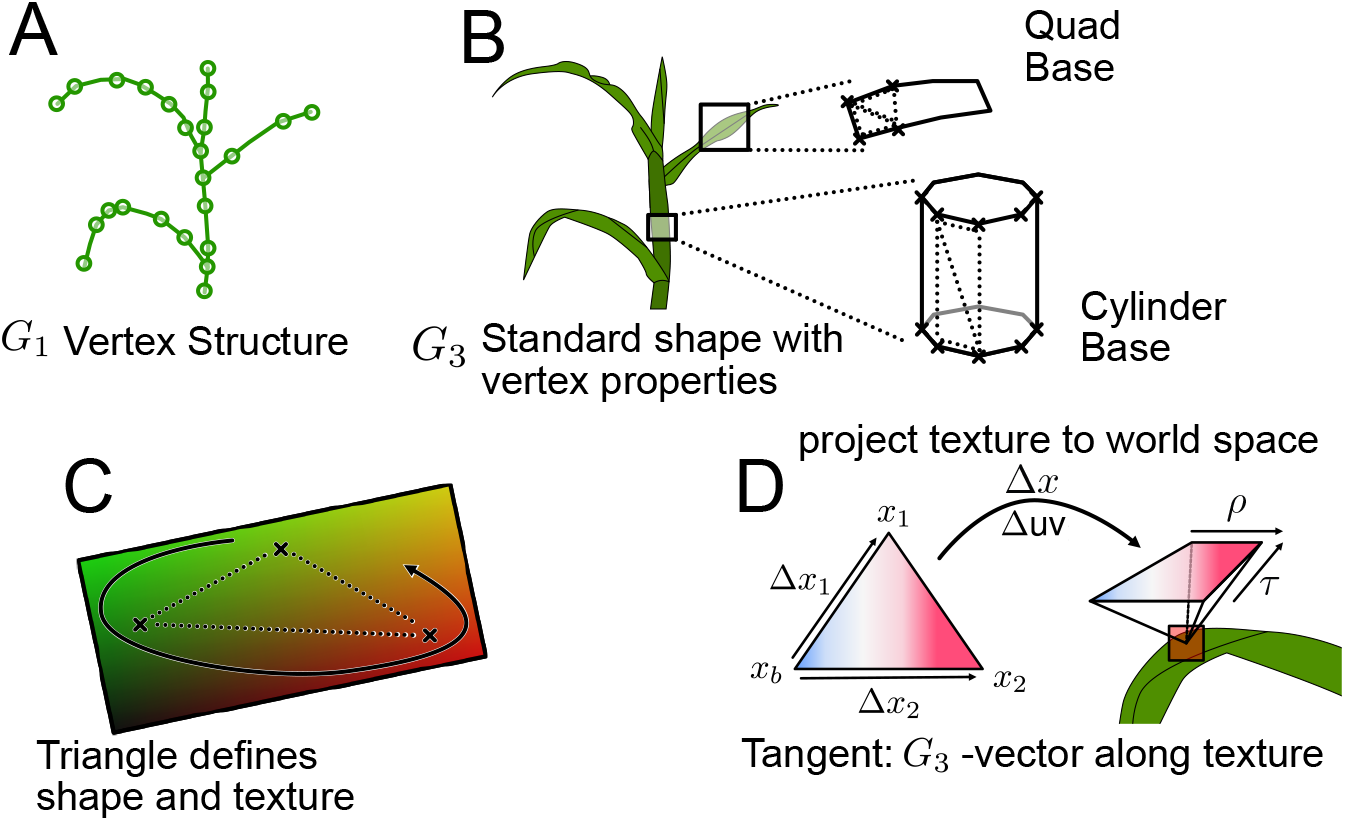
Relationship between functional properties on the texture level, according to the mapping employed by Synavis (Baker et al. 2023). **A**. We denote *G*_1_ to be the graph structure produced by the FSPM, including diameters, organ shape parameters, as well as functional aspects. **B**. Geometrization is done through shape extrusion, taking uniform shapes as template and placing them in a way that they encase the plant. Triangulization is the connection between the shapes that are used to build the geometry. **C**. On top of a triangle, the texture map is further defined on each vertex. **D**. To be able to fully map the texture onto the geometry, the tangents need to be computed, which indicate for the texture is wrapped locally around the geometry, being as long as one pixel is wide on that vertex.

### 2.2 CPlantBox Model Embedding

CPlantBox produces the structure of a plant, using a graph (*G*_1_ := (*V, E, P*)) that encodes one-dimensional properties along the edges. *G*_1_ is made of vertices (*V*) connected by edges (*E*) and stores a set of properties (*P*) defined on either the edges or the vertices. This structure can be seen outlined in Fig. 1.A. Our embedding typically uses an inferred 3D structure, called *G*_3_, which contains shape information informed by *G*_1_. *G*_3_ is a meshing based on shape representations (cylinder or quad) that is implicitly generated from vertex positions and organ type, as shown in Fig. 1.B. We make certain assumptions to define the output geometry, e.g., the leaf blades are perpendicular to the branching direction, and remain perpendicular to successive vertex differences. This can be calibrated empirically through the CPlantBox visualizer (Baker et al. 2023). This construction means that in most circumstances the properties on the surface triangles are defined through the edge in *G*_1_ they were constructed from. We interpolate properties to vertex level and implicitly further by encoding properties into textures, seen in Fig. 1.C, which can be accessed through tangent information like Fig. 1.D.

UE uses physics-based rendering, utilizing a bi-directional radiative transfer function (BRDF) *f* (*ω*_*i*_, *ω*_*j*_), which yields the outgoing light intensity for the angle *ω*_*j*_ according to the incident angle *ω*_*i*_. The function *f* is defined either through geometric or texture properties encoded in the Material shader that dictates how the surface reacts to light. A surface can also emit light, and the final color at the hit point *x*^*′*^ where the pixel ray intersects the surface will include both direct emission as well as reflectance from other surfaces’ emission (indirect light). The reflectance of a surface is represented semi-empirically via a single factor, representing implicitly the interaction of the light with the different layers of an object. As we are measuring the brightness, we are reversing certain aspects of the light simulation - e.g., a surface that would have absorbed a band of the light spectrum would now be brighter instead. This was implemented as such mostly for convenience, allowing the values to be representing of the actual absorption rather than its inverse. The BRDF is sampled across the incident angles of possible other light source contributions.

The sampling range is locally hemispheric, but the effective brightness is scaled with the surface normal. The surface normal *n* is interpolated from its vertex definition much like other properties (cf. Fig. 1.C). The space of surface angles Ω needs to be thoroughly sampled to gain a fully informed view of the scene brightness, also illustrated in Fig. 2. While there are certain optimizations implemented in Lumen, there are no intrinsic tricks that can be utilized to speed up the sampling, as it is entirely dependent on the geometric content of the scene. Texture properties rely on a sparse map UV : *G*_3_ *x* ↦ *p* ∈ ℝ^2^, illustrated on triangles of *G*_3_ in Fig. 1.C, showing potentially different scales between vertices and texture maps. As described in Baker et al. (2023), the texture mapping of the FSPM structure fills the space of the texture map, thus resulting in a stretching of the properties contributing to the transfer function. Since the texture mapping is done on vertex level, but the rendering will densely prompt for surface properties at every location, the evaluation of a specific point *x*^*′*^ relies on the texture map. Fig. 1.D shows how the texture (and thus functional) information is wrapped around the geometry, while the corresponding value retrieval for the texture is shown in Fig. 2. This estimation is linear, can be done before rendering (thus saving the tangents at vertex level) and allows for seamless stitching of a texture covering unconnected surfaces by explicit tangent declaration. The computation of a value from the functional properties at a specific surface location is described in Fig. 2. The primary surface properties^2^ that are implemented in UE that are of interest are diffuse, which is set to grey for the purpose of measurement, specular, roughness, and auxiliary properties such as subsurface, translucency, or masking. Evidently, our implementation uses strong heuristics that need continuous verification by full-spectrum models for radiative transfer. We break down the most relevant contributions to photosynthesis and light absorption to material properties. The diffuse property (set to (0.5, 0.5, 0.5)) is a measurement tool that does not contain any physical relevance. However, although we are using a single heuristic, one could separate three channels for the transfer model. The virtual world model that is being used here is shown in the flowchart in Fig. 4.

**Fig. 2:**
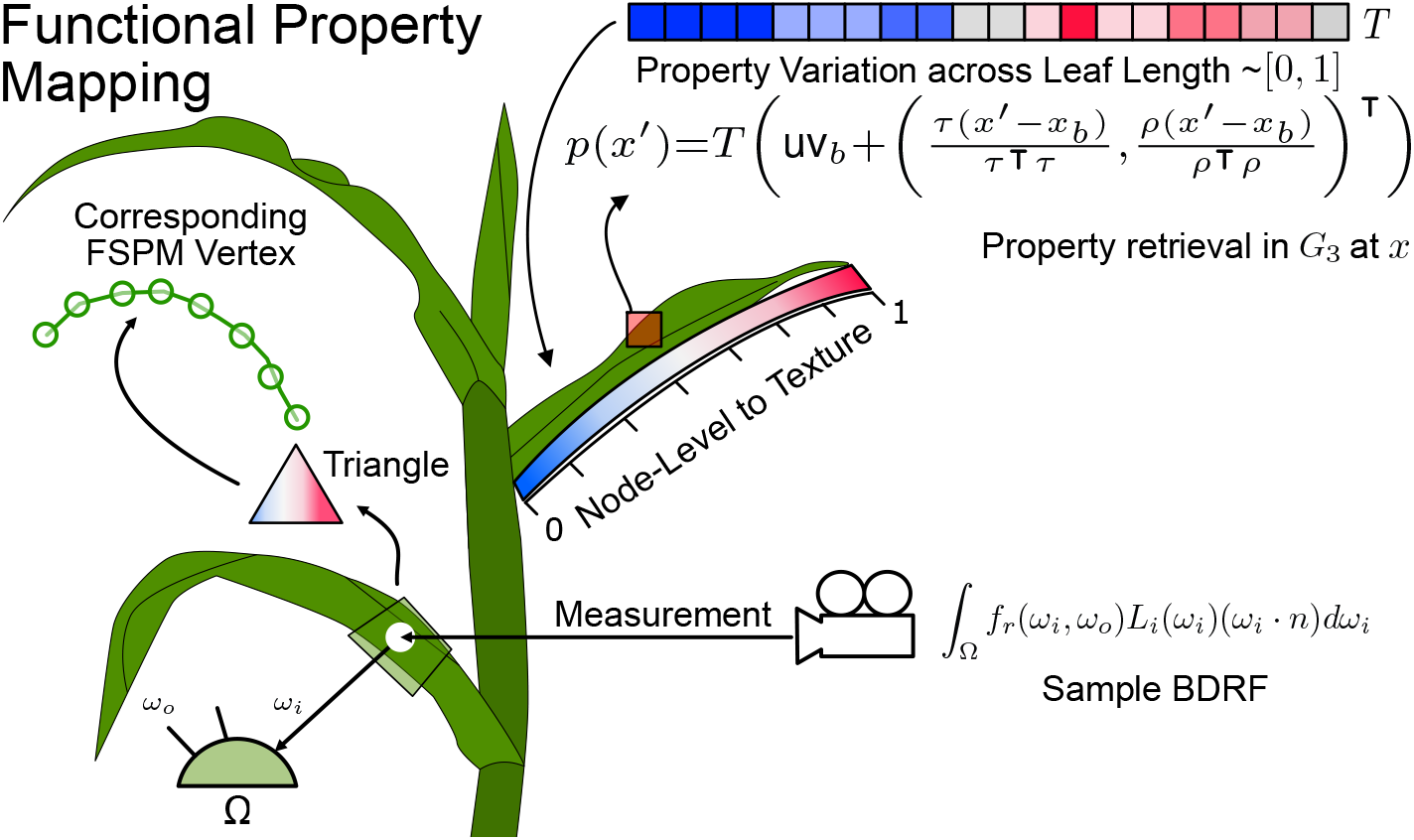
Matrix-notation description on how the functional properties are being sampled by the rendering pipeline, if defined through the Synavis framework in the prescribed way.

Emissive properties, such as chlorophyll fluorescence, are added to the total luminescence in the scene. This effect is smaller compared to the direct light influx in the scene, as characterized in Fig. 3. This characterization is primarily done to ascertain whether the impact of the emissive property can be calibrated. Surface light emission is being added outside of the hit-based sampling of the surface, meaning that the initial contact of the measurement ray yields a base luminescent contribution by the plant surface. In certain wavelengths of light, this behavior contributes to the total light absorption of the surface of the plants. As shown in Miao et al. (2018), this effect can have significant impact on the absorbed PAR rate (*mol/m*^2^*/s*) of the plant. Fig. 3 shows that the base emissive contribution will increase the base measured value, extinguishing in terms of impact with the increased directional intensity.

**Fig. 3:**
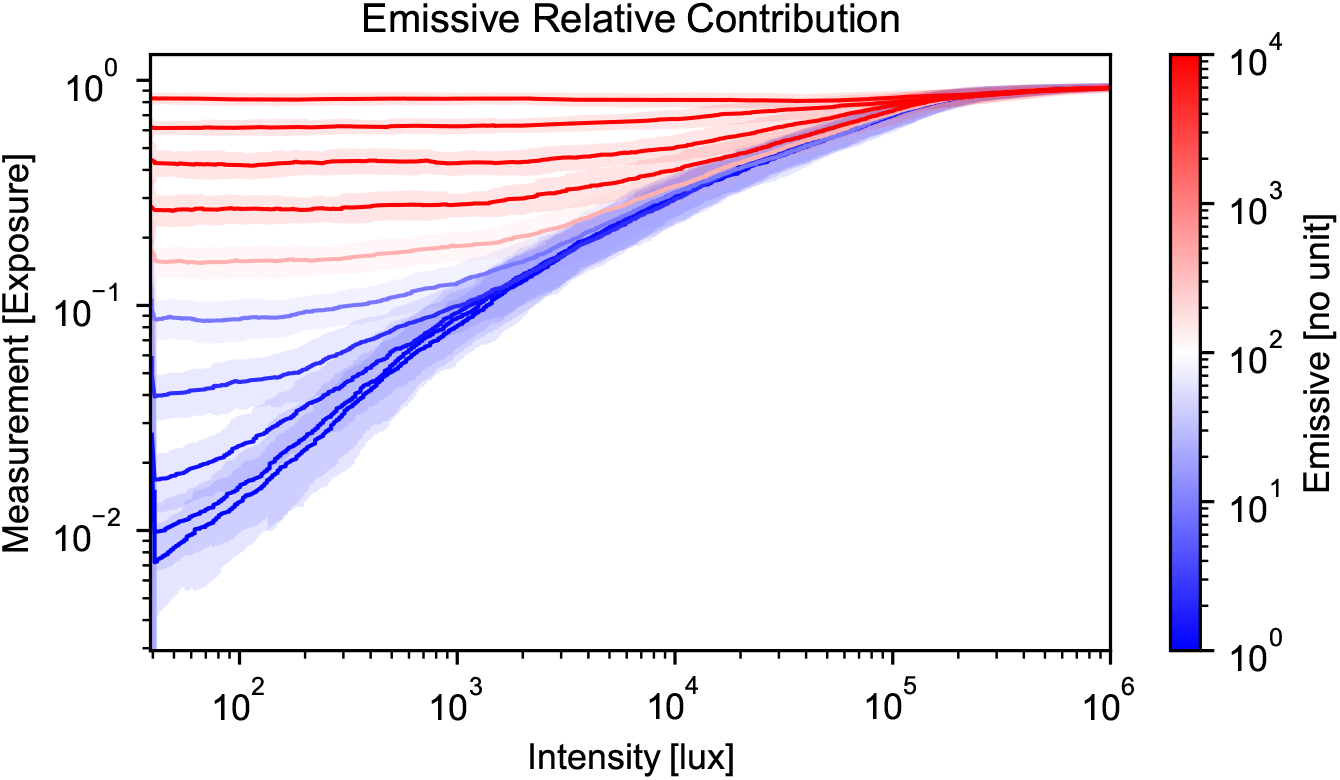
Characterization: relative contribution of emissive property on the BRDF rendering of the surface model in UE, depending on the brightness of the directional (sun) light. The measurement is non-scaled, meaning that the initial rendering system will output to a uniform [0, 1] range. On the x axis, the intensity is provided in *lux* units. We generally calibrate against an above-field sensor that is set to a direct exposure measurement reference.

### 2.3 Domain Partition and HPC

Since we are aiming to simulate field-scale setups, the actual workload of simulating and measuring in the virtual environment needs to be distributed. For this purpose, we utilize field partitioning by allocating instances of Unreal Engine dynamically through Synavis (Baker et al. 2024). Field distribution is based on direct partitioning. Because CPlantBox uses stochastic structures that are random-number-based, we assign a specific starting seed to the field in total, varying the individual plants deterministically. This ensures that the specific realization of a plant organ does not need to be communicated between nodes, as all nodes have implicit knowledge about the neighboring structures and could access their specific realization by simulating the corresponding field seed + plant position. All simulations that are being run have unique plant structures, but these structures can be inferred from the starting time of the cluster job.

We implement our field distribution for the JURECA-DC (Thörnig 2021) cluster, equiped with four NVIDIA A100 GPUs in addition to two AMD EPYC 7742 CPUs. These nodes are connected with InfiniBand Interfaces (NVIDIA Mellanox Connect-X6). Within each node, the field consists of a realization of a grid of plants, growing in a separate CPlantBox configuration. It is entirely optional, and independent of the virtual scene, whether the boundaries of the individual FSPM instances are being coupled, or not. In fact, as we are measuring the light influx on the geometry in the scene, the competition and individual exposure of individual plants can be estimated without running the FSPM in a continuous system. Nevertheless, for a full view of the plant field, a continuous system of plants, particularly regarding their soil properties, would be necessary.

The field-scale embedding further needs to be distributed over nodes to ensure efficient computation of plant instances. This is the case for all field-scale setups, irrespective of the soil-related simulation setup, since the amount of individual segments that need to be measured exceeds the memory of common GPUs (Baker et al. 2024). Our scaling experiment, results of which are shown in Sec. 3.3, consists of a partitioned field using a total number of plants *N* ^2^ arranged in a square. Based on total amount and MPI world size *W*, this square is subdivided into sets of squares of *N/W* plants per side. The seed numbers act as IDs for the plants, with the added randomness introduced by the starting time. The simulation can use one coherent global starting seed that is concatenated with the local ID of the plants. The grid of plants is scaled with the world embedding scale (which can be calibrated if needed for numerical precision) and the plant density. However, custom placements of plants are generally encouraged when producing light simulations, as our partitioning setup uses simple idioms to distribute the simulation as a showcase for HPC partitioning.

Our partitioning and scaling experiments are based on a plant grid that is being set and defined remotely. There are many hyperparameters that contribute to performance and results. In our tests, the total virtual screen resolution, i.e., number of light meters, is a factor in overall performance. For the hard scaling, we tested the progression using three steps of meter count: 50 ↦3200Pixels, 100 ↦6400Pixels, and 200 ↦12800Pixels. Virtual resolution values outside of these yield either decreased render or measurement performance. We increase the number of nodes, resulting in an increased field size or an increased distribution of the field size. In terms of UE rendering configuration, we do not utilize culling and will keep the reference point for render optimization^3^close to the target for measurement. When path tracing^4^is employed, lightmap settings do not matter, and we are not utilizing virtual textures^5^, though it would not impact the measurement if this was added.

### 2.4 Embedding of Experimental Data

Our reference data has been taken from the Field Minirhizotron Facility in Selhausen^6^ (Lärm et al. 2023; Nguyen et al. 2024). The facility is an experimental site that includes field data as well as extensive sensory data, including a minirhizotron setup that includes tubing to measure root distribution (Bauer et al. 2022). These data sets are hosted on the TERENO platform (Bogena 2016), which is an open data platform that enables us to access sensory as well as experimental data across parts of central Europe.

Weather is introduced both as environmental condition influencing the atmospheric boundary of the photosynthetic reaction and as precursor to the quantization of other parameters. From top to bottom, air pressure and humidity, temperature, and radiation are being derived from Selhausen measurements. These values are partially resolved at the 10-minute rate. Using the calibration in Fig. 2, we can linearly scale the light measurement using a clearsky measurement value as reference. This value can stem from singular measurements done as reference, or a densely resolved radiation flux measurement as present in the Selhausen above-ground data. From a data generation standpoint, PAR radiation can be directly mapped onto the pixel space by using the relative radiance at each point without needing to consult measurement data. An illustration of this approach is shown in Fig. 4. We vary the directional light brightness based on the RadiationGlobal [*W/m*^2^] data collected in the Selhausen facility, as illustrated in Fig. 4. Our site data was partially extracted from the TERENO data hub (Bogena 2016) for the climate station Selhausen 3 (Schmidt 2024). The soil characterization was done in Lärm et al. (2023), which features matric potential measurements at different depths. The soil parameters were implemented as one-dimensional model for simplicity, while the root structure was simulated explicitly. We chose not to infer air co_2_ molar fraction from the experimental measurements as the sensors we would have available are not representative of the chamber measurements in Nguyen et al. (2024). Nevertheless, the full weather model can be referenced to in the simulation script, see Sec. 5.4.

For the 2016 measurement, winter wheat was sown on October 26th, 2015, while first emergence was logged at November 1st, 2015. Nguyen et al. (2024) describe the gas chamber measurement in the field regarding the photosynthetic activity of the plants. The measurements were done at multiple time of days, and the PAR was always measured concurrently, along with other indicators. Our comparison focuses on the PAR adsorption and the co_2_ uptake. We assumed the start of the growing season to be 5th March 2016 and simulate the plant growth up to the measurement, at which time we run the photosynthesis module in combination with the UE measurement of light influx. We are using a pre-flowering gas chamber measurement for the photosynthesis activity, including a sunlit wheat plant. We measure the light photon influx per segment on the leaf surface. This value has been relayed to CPlantBox, following a photosynthesis evaluation with simplistic soil coupling. We assumed no limitation by soil water content. The leaf surface parameters are kept simple, not including subsurface scattering or fluorescence. These values are included in the model, but need to be parameterized, and our TERENO-based parametrization is illustrated in Fig. 4. Both the virtual world models are steered through Synavis and implemented in UE, while CPlantBox runs as a precursor to the data scene, needing the model scene as light influx reference. In UE, the measurement was done over 3 frames following a 2 frame waiting period after the measurement point has been registered. This is to ensure that the virtual scene is fully updated before the measurement begins.

**Fig. 4:**
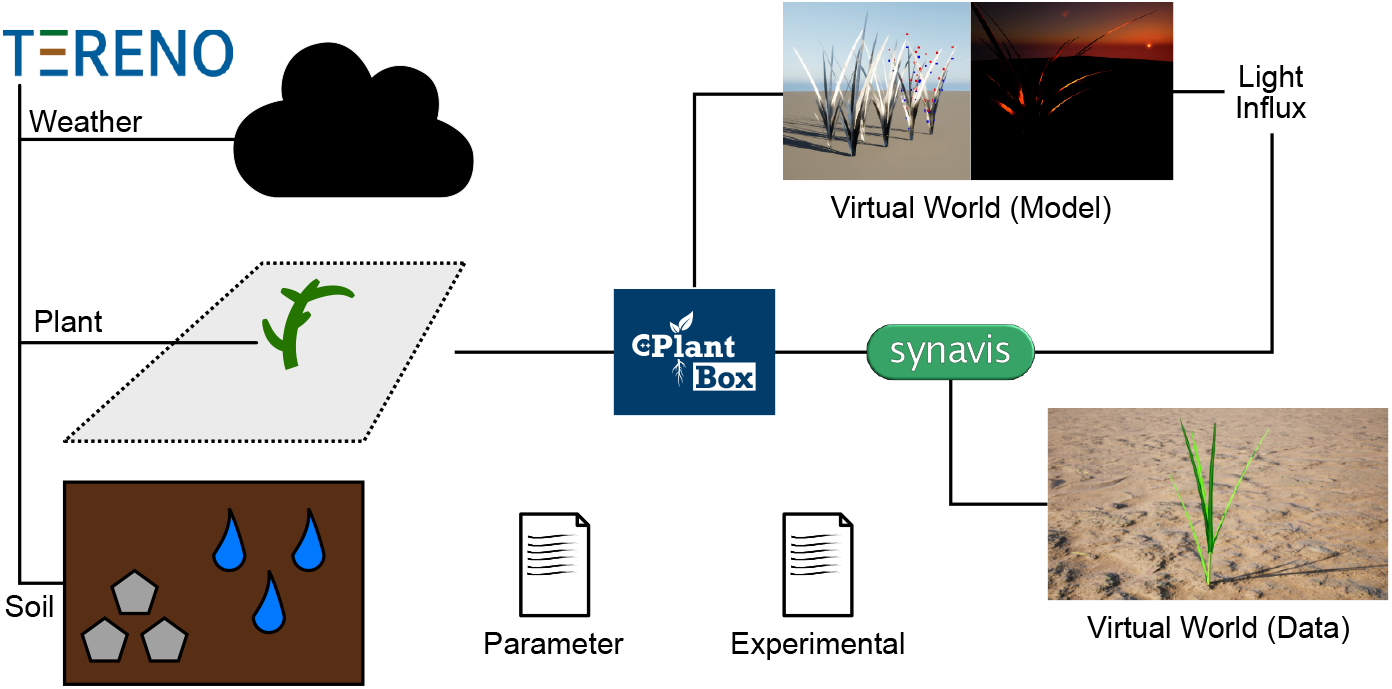
Data sources and pathways of the UE virtual world coupling for photosynthetic radiation influx measurement. CPlantBox is the central data source for UE, while on the left are static data sources. The model world provides light influx, which can be mapped onto the data world.

## 3 Results

Model coupling is largely based on the combination of the virtual world embedding (Baker et al. 2023) and the photosynthesis calculation developed by Giraud et al. (2023). Here, we highlight our results regarding the primary evaluation of our coupling’s main contributions. This section focuses on the accuracy of the framework, the scalability of its implementation, as well as the production of synthetic light exposure training data from FSPM simulations.

### 3.1 Synthetic Light Exposure Data

Mapping the functional data onto a virtual measurement space ensures the increased usefulness of the photosynthesis data. It allows the extension of the synthetic data pipeline to uncover mechanistic insights into plant development in a pipeline otherwise restricted to other information that is being rendered in the engine, such as implemented by Mousavi et al. (2020).

Our UE implementation measures exactly around the submitted points, an array which can be mapped onto the geometry. As the rendering needs to switch modes for this task, the workload needs to be separated from the measurement or performed subsequently. The photosynthesis module of CPlantBox, if not restricted through other mechanisms, imposes a limitation on carbon uptake *A*_*g*_, which is *A*_*g*_ = min (*F*_*j*_, *F*_*c*_) (Giraud et al. 2023), where *F*_*j*_ is the electron transport rate and *F*_*c*_ is the carboxylation rate, which is largely restricted by Nitrogen uptake. These rates are output by the simulation, particularly the photosynthesis module, when solving for the photosynthesis reactions. They are given by segment, which we interpolated to organ level for the visualization in Fig. 5. Different populations, one large and one rather small are shown in Fig. 5, showcasing how this kind of scaling would function in the Synavis pipeline. An output of this simulation is the fact that we can scale such an otherwise very mechanical value to a more clearer result, showing a potential limitation by *F*_*c*_ which could indicate the need for more nitrogen fertilization.

**Fig. 5:**
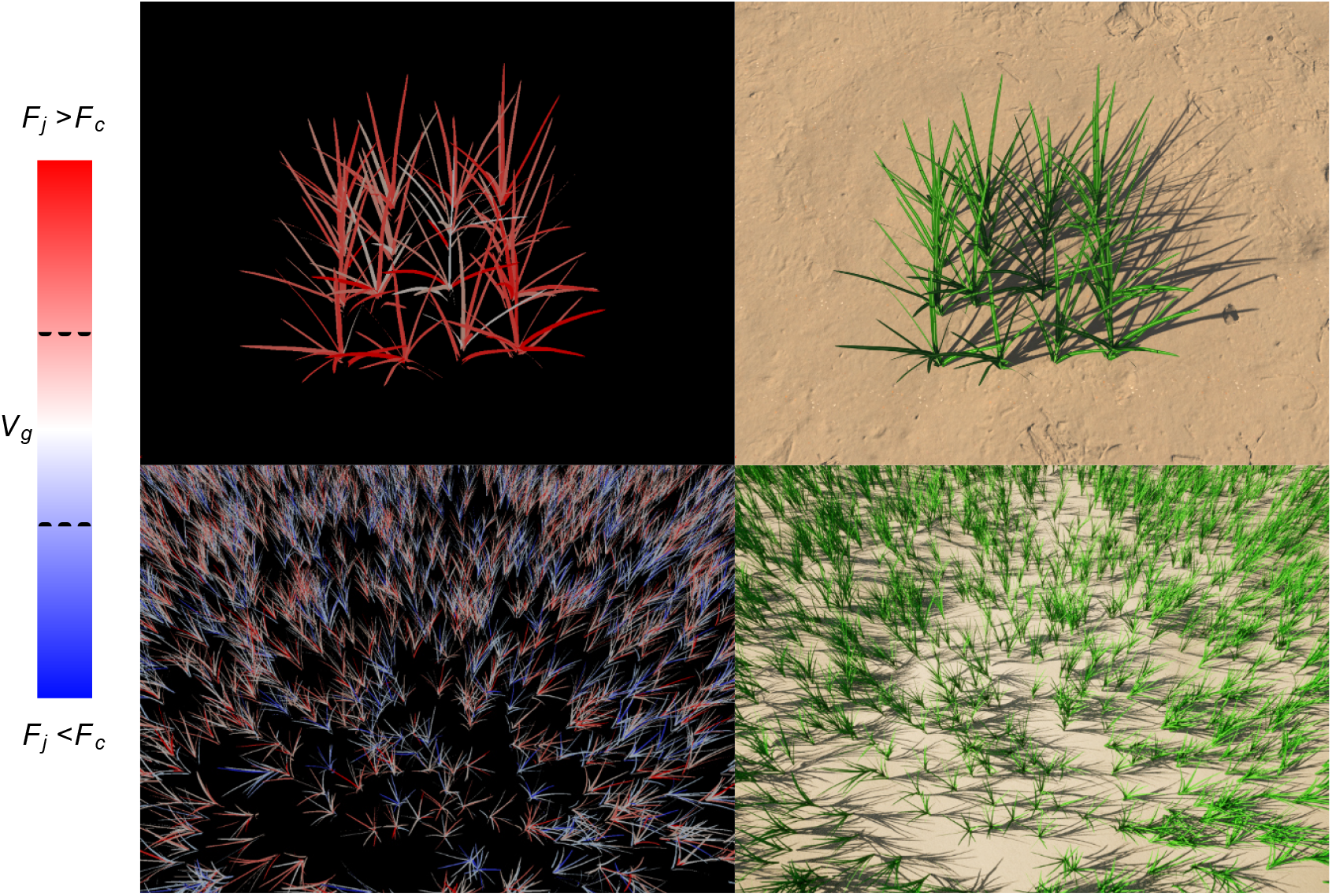
Mapping of functional properties, in this case light assimilation effectiveness, onto the leaf area of the FSPM simulation. These values are scaled mostly categorical, with the red/blue distinction describing the flipping point between nitrogen limitation *A*_*n*_ and photoelectric limitation *V*_*j*_.

Our implementation for this work furthermore allows a full export of the FSPM geometry for multiple file formats, including VTK-compatible files, or Wavefront Object files that can be used with software such as DART (J. P. Gastellu-Etchegorry and Gascon 2004). The higher compatibility also allows users to employ more coupling techniques that can be used with other RT models, including lightweight models that run on CPU-only devices. The coupling is furthermore entirely steerable within Python, which also contains the simulation information provided by CPlantBox.

### 3.2 Light Influx Accuracy

We compare the experimentally measured values for light influx and carbon uptake to our virtual world model. Our comparison is shown in Fig. 6, which highlights both light influx as well as co_2_ uptake per area. Our calibration yielded a close match to the experimental data, though we will note that multiple parameters could yield similar results. The calibration of the individual parameters used for the light evaluation can be referenced to in the resources that accompany this manuscript, see Sec. 5.4.

**Fig. 6:**
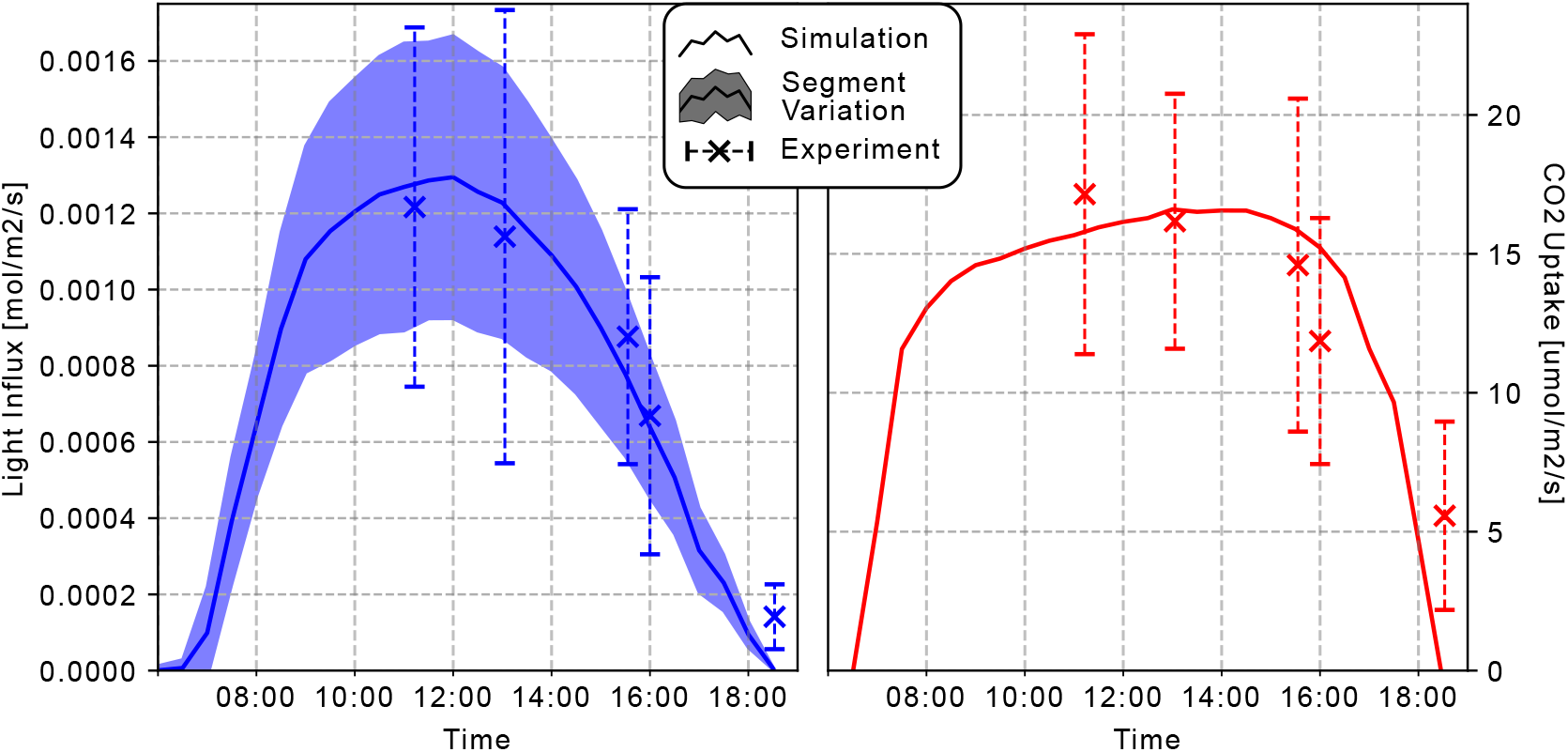
Absorbed PAR in blue/left, and net co_2_ uptake in red/right. The light flux standard deviation is segment based, i.e., depending on where on the plant the light is measured, the light measurement has different results, resulting in a high standard variation. The experimental data is marked with an x and a dotted line for measured variance.

For our simulation study, we sub-sampled the 10-minute intervals to 30-minute intervals to reduce simulation time to allow for a better progression of the FSPM simulation. This is mostly a reduction of the computational workload to replicate the data, while still maintaining similar expressiveness. The base experimental time resolution of the experiment is non-uniform, with the measurements taking place in specific time slots with multiple measurements. We averaged measurements that were taken in the same interval, though on different plants labeled the same, resulting in the variance seen in Fig. 6. Similarly we highlight that while the measurements that were deemed *sunlit* in the experiments described in Nguyen et al. (2024). The light absorption measured in both the experiment and in our simulation study yielded high variance. This is likely caused by differences in measurement positions on the plants. The increase in the mornings was steeper than the decrease in the evening. This is also present in the raw data we used for the modeling (Schmidt 2024). Similarly, the co_2_ uptake increased in such a fashion, though in the evening both radiation and uptake values are outside the experimental range. Our simulated co_2_ uptake at around 11 ó clock was lower than the average experimental value, though within the variance. Our light influx variance was peculiarly low in the range of 7 to 8 ó clock, which we could attribute to the scattering values that were used in the virtual environment, in combination with the fact that overall illumination was low.

### 3.3 Scaling and Performance

We have investigated the scalability of the coupling to determine the performance of the method on HPC systems. We show both timings and resulting speedup in Fig. 7. In it, we observe what appears to be super-scaling behavior, which requires closer investigation. First, the light evaluation being distributed across nodes will make the measurement faster than on a single node, with *N/P* measurement points as opposed to *N* . We note that there *is* overhead in the shape of the surrounding plants that get rendered as to not distort the boundaries between the compartments of the field, even though these plants act as geometric inclusion only. However, because the overall amount of triangles in the virtual scene is significantly lower, the engine also performs better *per render pass*, which results in an increased performance of a single evaluation. Thus, the parallel performance appears to be so steeply increasing with the number of nodes. This is commonly referred to as memory-bound performance (see e.g., Allande et al. (2014)).

**Fig. 7:**
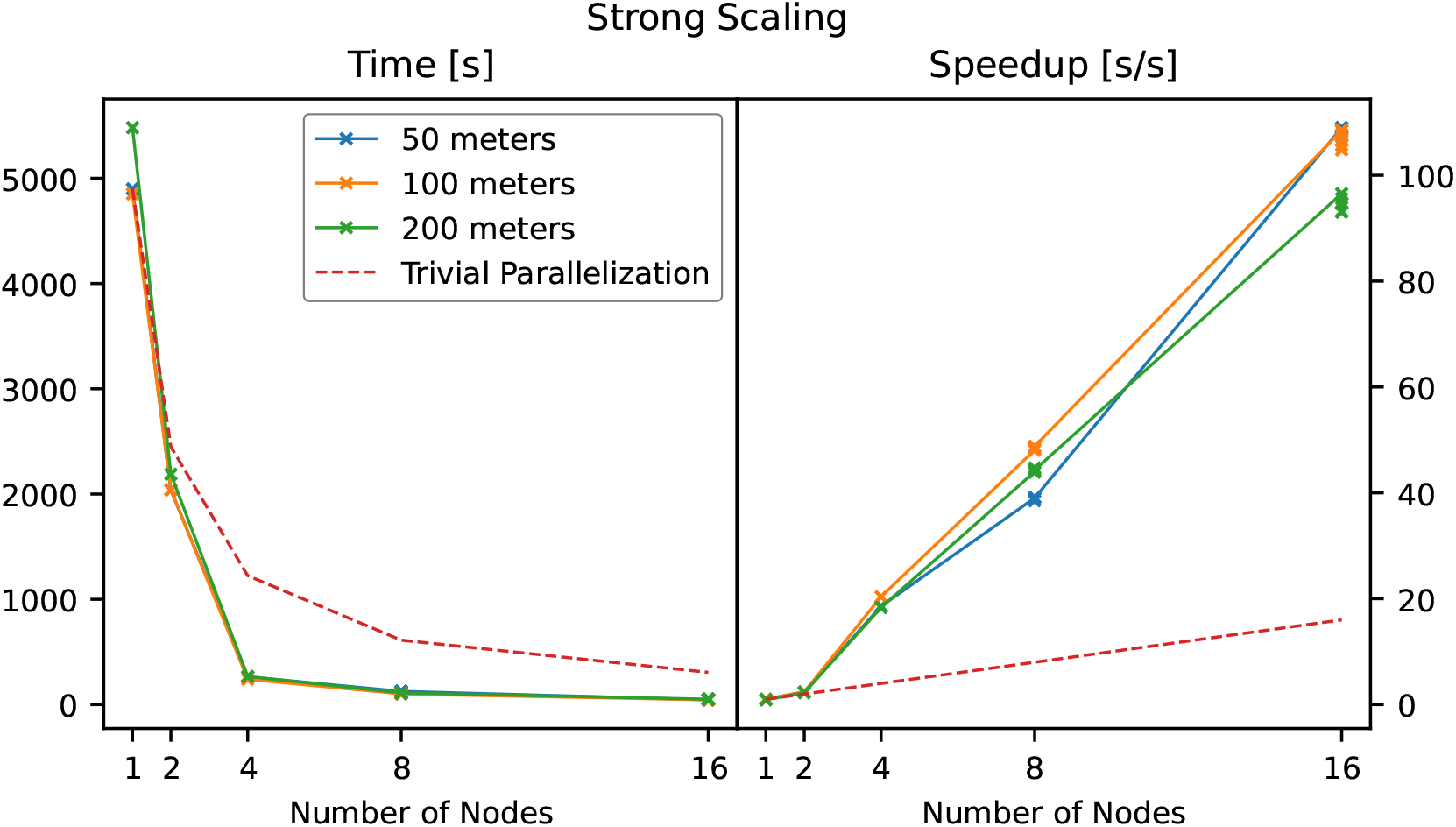
Scaling of number of nodes while not scaling total problem size. A: Time of execution for the photosynthesis pipeline for a field of 500 plants, measuring the performance of the light evaluation only, from submission to reception of the exposure data. B: Derived speedup values for the parallel performance. We compare our results against the ‘trivial parallelization’ baseline, representing a speedup directly proportional to the number of nodes used. Our memory-bound applications has a superscaling behavior, resulting in an overall faster computation by means of parallelization and reduction of wait cycles for the computation.

Fig 8 shows the weak scaling of our system. The speedup appears to be slightly increasing, though this is constraint by the controlling node that handles the same area. From measurement, the added speedup even though the problem size remains constant might have different reasons, as investigated in Sec. 4.1. The measurement is taken through the framework itself, and as such it does not account for effects due to MPI setup and the initialization phase of the photosynthesis simulation. In our evaluation, we found that there was not much difference in the performance regarding the total (virtual) resolution of the light measurement, based on the top three performing settings in this context. An increased resolution, i.e., an increased number of light meters, will result in a decreased frame performance, resulting in a relatively similar performance overall. From the hard scaling comparison in Fig. 7, we found that 100 light meters yields best performance, resulting in a total resolution of 8^2^ · 100 = 6400 measurement points.

**Fig. 8:**
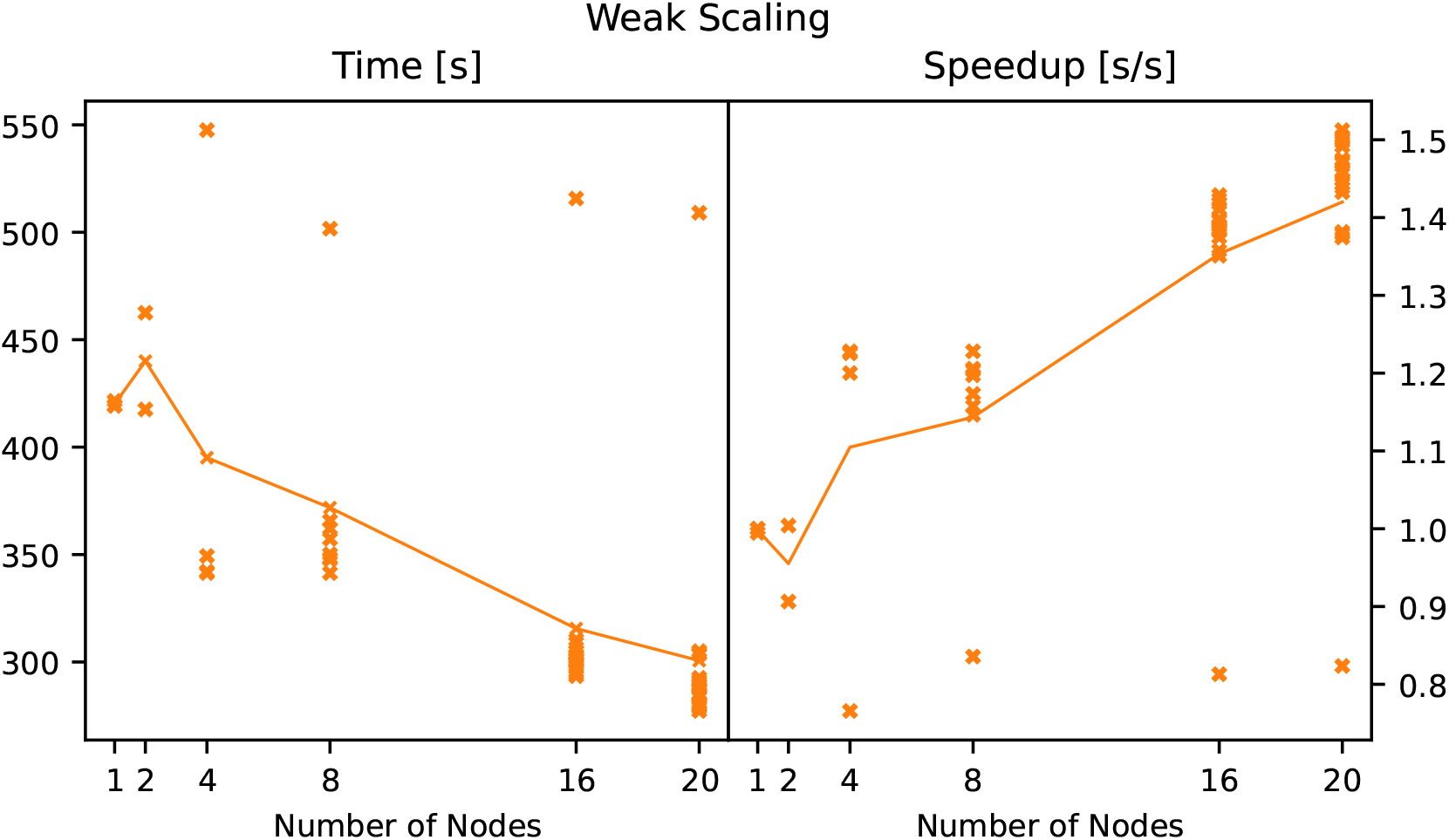
Scaling of both problem size and number of nodes. A: Time of execution if we increase the data in the same way as the number of nodes. B: Derived speedup values for the parallel performance, again calculated as *T* (1)*/T* (*N*).

## 4 Discussion

This work has been an investigation into the validity of using the Synavis framework for simulated embeddings, lifting the data production pipeline to a true embedding of simulated data. The contribution on a technical level is the coupling to the FSPM as well as the implementation of a functional embedding into UE such that it can be used together with other models beyond plant models.

### 4.1 Distributed Simulation of FSPM Embeddings

It is generally no issue to distribute plant simulations across nodes in cases where the simulation is entirely at organ scale. Scale conversions, such as between cells and plant level, always require a form of heuristics, as it would be cumbersome to simulate a plant on cell level entirely. As such, the data scale conversion is an important tool to not only use varying resolutions of experimental data, but also to distribute the plant simulation. We use this to estimate the size of the buffer region in our tests. Typically, with our parameters regarding the size and density of the field, two rows of plants around our measurement area is sufficient on average. During midday, and depending on geographic location, the buffer plants might even be omitted, which is also the case for cloudy days. In the first case, this is because the shadows are very close to the plant and might not significantly touch neighboring plants, and in the second case, most light arrives in so many scattered pathways that shadows have decreased contrast.

The scaling experiments in Fig. 7 and Fig. 8 show a clear speedup through HPC use. We note, however, that we do not believe the apparent trend in Fig. 8 regarding the speedup between 8 and 20 GPU nodes has a systematic cause. There are components regarding the order of operations, particularly the spawning of plants and the initialization of the measurement, that cause a greater variance in the sampling. Particularly we note that the main compute node did not speedup, but rather remained somewhat constant regarding its sampling and the collection of light influxes. Uncovering the hyperparameters that contribute to the eventual parallel efficiency is beyond the scope of this work, but would be the next step to complete the HPC integration of the Synavis framework, particularly in combination with AI workloads. To accommodate DNN pipelines and to reduce over-fitting, the number of visualizers should be lower than the number of learners, as the learning algorithms needs to use image-based augmentations in addition to domain-based augmentations to fully make use of the synthetic data, as image-based augmentations have uses beyond adding more pictures to the data space.

Distributed performance using the Unreal Engine also would have some limitations regarding inter-module connectivity, as scaling up the virtual world rendering on a dedicated visualization module might impact the latency. Users of the framework would need to be cognizant of the local properties of the HPC systems they operate on, as it could introduce topology or moduledepended differences in communication times. Our communication tests typically yielded latencies between modules (through Ethernet, e.g. between JURECA-DC and other clusters on-site) of around 200ms. We do highlight the usefulness enabling graphics on user-exclusive GPU nodes as opposed to offloading the capability of rendering onto interactive nodes that cannot tolerate high workloads.

### 4.2 In-Silico Recreation of a Field Experiment

The gas chamber measurements can be referred to in Nguyen et al. (2024), and we will note that it is not an isolated chamber, and thus the concentrations are susceptible to wind. We are comparing distribution-free measurements, as the reported photosynthesis measurements are provided per area and time. Our pipeline is able to replicate the experiments of this specific configuration well. We have to note that there are many configurations of parameters that might yield similar results, some of which are outside what are realistic properties for plants. The overfitting of simulation models has been previously reported in other domains, such as radiotherapy in silico simulation (van der Schaaf et al. 2012). The calibration of the light influx is not only dependent on the actual sunlight intensity, but also on the atmospheric scattering of specific wavelengths, as well as leaf surface scattering. To truly provide an embedding for more different FSPM simulations, a full study, including experimental measurements of leaf surfaces, would be required, along with a study on the mapping of wavelengths into the virtual world. We have fitted and highlighted that we can represent experimental data, but the full parametrization and analysis of the virtual world is beyond the scope of this paper, particularly regarding its use as data generation pipeline. The soil parametrization regarding light propagation is an important factor that contributes to total illumination.

The experimental replication furthermore yields a type of bottleneck that correlates with the production of synthetic data. In future work, less light influx settings, i.e. day, time, and weather, should be computed to speedup evaluation, in addition to only computing a representative subset of the field. This would speedup the total evaluation significantly, and more time can be allocated for the actual data generation, after deriving a heuristic of the mapping between the parameter space and the light influx or carbon uptake limitation on the plant. For the domain composition, the challenge poses itself that a camera either needs to be facing downwards, as this is how the light evaluation is parallelized, or the render scene needs to be setup by reducing the decomposed scene into one.

### 4.3 Functional Synthetic Data using UE

One factor contributing to the increased use of UE in certain domains, specifically in plant science, is the ability to very quickly prototype scenes. The key aspect is that visually, a user of these engines can achieve a good general setting very quickly, beyond the need for texturing and geometrization, which are issues that are largely independent of the concrete rendering back-end. Examples of this are Helios or DART with subsequent rendering, or any alternative to UE for synthetic data implementation. These models also require calibration of surface properties, even if respective rendering pipeline use multi-spectral or physics-based models.

There are certain aspects of the pipeline that required adjustment seeing as UE is historically a game engine, which goes beyond terminology. Certain parts of our workflow are implemented on top of basic optimizations we explicitly turn off, or circumvent using procedural geometry. Similarly, certain features might not be as fleshed out as others, particularly regarding the implementation of simulation models, requiring some fine-tuning in cases that extend beyond what a game engine typically delivers. In Sec. 3.2 we showcase the connection between the simulation and the eventual training output that could be used for inference in field settings. There are caveats, but generally, our method adds to the virtual scene in a graphics engine information that is otherwise not available, by employing simulation models to act as data source.

While this work focuses on the central validity and applicability of the workflow for photosynthesis problems, this pipeline is a generic data production pipeline. More functional information within the virtual world would enable a more informed embedding and ultimately strengthen the ability of deep neural networks to estimate plant health through remote sensing techniques. Camera footage together with distance sensors are comparatively low-cost and easily reproducible in experiments, and they are also always computed in UE, as pixel shaders depend on pixel depth and relative velocity information. Camera settings, particularly within the data production, but also for the real world, are extremely important. In Fig. 5 we showcase a functional mapping and visualize the plants in a way that makes them easily distinguishable for this manuscript, but the scene is technically overlit, as two pixels yielding different light absorption information might have the same scene color value. This is an issue that is a vital component of any functional analysis that depends on consumer camera images. Unless the exposure is configured to take only leaf surfaces into account, there will be some loss of information due to the restricted brightness range of the images. This is the case even in UE, which internally uses a high dynamic range system but is flattened into pictures to produce synthetic data. We have previously reported the dependency of threshold-based relative leaf area on image capturing conditions in Baker et al. (2023).

## 5 Conclusion

This paper presents a coupling approach that is directly embedded into a data generation framework. Recent advances in data generation, particularly for agricultural science, have highlighted the need for embedded models in these frameworks. Here, we proposed a coupling with an FSPM model, highlighting the usefulness of its functional simulation for the enrichment of the data generation. A new implementation of radiative transfer representation in UE was developed to couple the FSPM CPlantBox to a light influx evaluation for photosynthesis evaluation. We show that we can embed functional properties into the virtual scene accurately, scalable, and usable for synthetic data.

We have included a replication of an experiment, a data production pipeline for light exposure efficiency and the carbon uptake limitation in plants, as well as the scalability of our method on HPC systems. Exploring how to best utilize the tools available, especially when implementing the simulations into a wider workflow, is essential.

## 5.1 Author Contributions

This work was written, revised, and edited by all authors. DB is primary author and conducted the implementation, data analysis, visualization, and drafted this work. DB,MG,JHG adapted the Synavis/UE software for this work. MR,JHG provided access to supercomputing resources. MG,HS,AS provided domain expertise and guided the implementation of the plant simulation and experimental replication. EH,AS guided the data analysis and research aims of this work. MR,EH are the primary supervisors for the University of Iceland. AS is primary supervisor of this work.

## 5.2 Acknowledgments

The authors thank Thuy Huu Nguyen for his advise on data interpretation and use. Furthermore, the authors acknowledge compute time granted to the JURECA-DC Supercomputer in the project visforai.

The authors would like to acknowledge funding provided by the German government to the Gauss Centre for Supercomputing via the InHPC-DE project (01H17001). This work has partly been funded by the EUROCC2 project funded by the European High-Performance Computing Joint Undertaking (JU) and EU/EEA states under grant agreement No 101101903. This work has partly been funded by the German Research Foundation under Germany’s Excellence Strategy, EXC-2070 - 390732324 - PhenoRob and by the German Federal Ministry of Education and Research (BMBF) in the framework of the funding initiative Plant roots and soil ecosystems, significance of the rhizosphere for the bio-economy (Rhizo4Bio), subproject CROP (ref. FKZ 031B0909A).

## 5.3 Conflicts of Interest

The author declare that there is no conflict of interest regarding the publication of this article.

### 5.4 Data Availability

The Selhausen data sets are described in Lärm et al. 2023 and Nguyen et al. 2024, and all relevant download information can be seen in those articles. Our framework, Synavis, is Open Source and the relevant codes can be seen in dhelmrich/synavis. CPlantBox is open source and available at Plant-Root-Soil-Interactions-Modelling/CPlantBox.

A video description is available at 10.6084/m9.figshare.28280780. The simulation code that is associated with this manuscript is available in the feature/experiment branch and is scheduled to be merged.

## Footnotes

1 Unreal Engine Source Documentation, account needed for source code access

2 Unreal Engine official documentation on the topic of Physically Based Materials, Accessed February 6, 2025

3 Occlusion Culling Description

4 Path Tracing Description

5 Virtual Texturing Description

6 50.865°N, 6.447°E, 203m a.s.l. in 2024

